# H_2_O_2_/ABA signal pathway participates in the regulation of stomata opening of cucumber leaves under salt stress by putrescine

**DOI:** 10.1101/2020.08.28.272120

**Authors:** Siguang Ma, Mohammad Shah Jahan, Shirong Guo, Mimi Tian, Ranran Zhou, Hongyuan Liu, Bingjie Feng, Sheng Shu

## Abstract

The stomatal-aperture is imperative for plant physiological metabolism. The function of polyamines (PAs) in stomatal regulation under stress environment largely remains elucidate. Herein, we investigated the regulatory mechanism of exogenous putrescine (Put) on the stomatal opening of cucumber leaves under salt stress. The results revealed that Put relieved the salt-induced photosynthetic inhibition of cucumber leaves by regulating stomatal-apertures. Put application increased hydrogen peroxide (H_2_O_2_) and decreased abscisic acid (ABA) content in leaves under salt stress. The inhibitors of diamine oxidase (DAO), polyamine oxidase (PAO), nicotinamide adenine dinucleotide phosphate oxidase (NADPH) are AG, 1,8-DO and DPI, respectively and pre-treatment with these inhibitors up-regulated key gene NCED of ABA synthase and down-regulated key gene GSHS of reduced glutathione (GSH) synthase. The content of H_2_O_2_ and GSH were decreased and ABA content was increased and its influenced trend is AG>1,8-DO>DPI. Moreover, the Put induced down-regulation of ABA content under salt stress blocked by treatment with H_2_O_2_ scavenger (DMTU) and GSH scavenger (CNDB). Additionally, the application of DMTU also blocked the increase of GSH content. Collectively, these results suggest that Put can regulate GSH content by promoting H_2_O_2_ generation through polyamine metabolic pathway, which inhibits ABA accumulation to achieve stomatal regulation under salt stress.

**Highlight:** Exogenous putrescine alleviates photosynthesis inhibition in salt-stressed cucumber seedlings by regulating stomatal-aperture.

## Introduction

In order to meet the growing food demand, it’s urgent to develop a stress resistance adaptive mechanism, which helps to increase crop yield (Fahad *et al*., 2015). Carbon dioxide absorption capacity of the most plant leaves become lower due to the salt stress (Elbasan *et al*., 2020), and has a significant effect on net photosynthetic rate (P_*n*_), stomatal conductance (G_*s*_), transpiration rate (T_*r*_) and intercellular carbon dioxide concentration (C_*i*_) (Moles *et al*., 2016; Elbasan *et al*., 2020). If C_*i*_ decreases with the decrease of G_*s*_, the decrease of P_*n*_ is caused by stomatal limitation, otherwise, it is caused by non-stomatal limitation. Therefore, the effects of salt stress on plants can be divided into stomatal limitation period and non-stomatal limitation period (Hejnak *et al*., 2016; Xia *et al*., 2017; Yang *et al*., 2019). At the early stage of salt stress, stomatal closure is the main limiting factor (Sudhir *et al*., 2004; Yildiztugay *et al*., 2020) and decrease the activity of ribose 1,5-diphosphate carboxylase/oxygenase (Rubisco), destroy the photosynthetic system, and the chloroplast structure/function or thylakoid energy over excitation are the main limiting factors, at the later stage of salt stress (Chen *et al*., 2020; Sudhir *et al*., 2004; Demetriou *et al*., 2007; Shu *et al*., 2012). In turn, P_*n*_ function inhibited and carbohydrate metabolism becomes destroyed, thus reducing dry matter accumulation (Jurczyk *et al*., 2016). Therefore, it is essential to increase stomatal pore size during stomatal limitation period to resist salt stress injury.

In recent years, a large number of research reports on the role of PAs and ABA in resistance of higher plants to salt stress has been published. Putrescine (Put), spermidine (Spd) and spermine (Spm) are the most abundant PAs in plants, which are state in the free, conjugated and bound form in plants (Pál *et al*., 2015). One of the main functions of PAs is to balance the ROS (reactive oxygen species) level that induced by NaCl stress and provide a suitable physiological environment for cell survival and metabolism (Saha *et al*., 2015; Seo *et al*., 2019). Exogenous Put could effectively improve G_*s*_ and P_*n*_ of cucumber leaves under salt stress (Zhang *et al*., 2009). In addition, PAs may also act as stress messengers in plant response to different stress signals, as a regulatory signal molecule, induce different transcriptional responses (Liu *et al*., 2007; DU *et al*., 2019; Marco *et al*., 2011). ABA, as a plant hormone, plays an important role in the adaptive response to abiotic stresses that cause cell water loss (Zheng *et al*., 2019). Since, G_*s*_ is largely controlled by ABA, the change of ABA concentration is often used to explain the change of stomatal morphology (Verslues *et al*., 2006). Studies have shown that PAs participates in signal transduction through the interaction with other hormones (Bitrián *et al*., 2012), such as ABA (Alert *et al*., 2011; Zhu *et al*., 2020). Espasandin *et al*. (2014) found that biochemical and morphological responses of cell water loss were related to Put level and Put regulated ABA biosynthesis at the transcriptional level to control ABA content in response to drought, indicating that these two plant growth regulators have complex cross-sectional effects in stress response. However, it is still unclear how to Put interacts with ABA to regulate stomatal changes during stomatal limitation period under salt stress and hence, we conducted the current experiment to extends our knowledge about this regulation. H_2_O_2_ is produced by the catalysis of PAs catabolism enzyme DAO and PAO, which are located in the apoplast and plays an important role in plant defense against abiotic stresses (Alcázar *et al*., 2010; Tavladoraki *et al*., 2012; Guo *et al*., 2014; Fraudentali *et al*., 2019). The other two enzymes that catalyze the production of H_2_O_2_ are NOX and CWPOD (Rejeb *et al*., 2015; Passardi *et al*., 2004). It is noteworthy that GSH acts downstream of H_2_O_2_ in a number of processes, and GSH has been shown to negatively regulates ABA-induced stomatal closure (Ogawa, 2005; Okuma *et al*., 2011; Akter *et al*., 2013; Asgher *et al*., 2020). Therefore, it is imperative to solve whether and how H_2_O_2_ and GSH play a role in PAs induced stomatal response as well as find out the crosstalk between PAs and ABA.

We mainly explored the positive regulatory effect of exogenous Put on cucumber leaf stomata under salt stress and speculated the interaction relationship among the Put, ABA, H_2_O_2_ and GSH, and finally concluded the internal mechanism on how exogenous Put improving cucumber photosynthetic capacity under salt stress.

## Materials and methods

### Plant material and treatments

Cucumber variety “Jinyou 4” was used in this experiment, which was provided by Tianjin Kerun Research Institute, China. Uniform seeds were soaked in distilled water followed by placed into a constant temperature shaker (28 ± 1°C, 200 rpm) for germination. After 24 h, germinated seeds were sown in a 32 hole seedling tray filled with quartz substrates, and cultured in Solar Greenhouse at Nanjing Agricultural University, where the diurnal temperature 28 ± 1°C/19 ± 1°C (day/night), and the relative humidity 60-75% were maintained. After second true leaves were fully expanded, seedlings were transplanted into plastic containers for pre-culture with half-strength nutrient solution (pH 6.5 ± 0.1, EC 2.0-2.2 mS·cm^−1^), and air pump was used to stabilize dissolved oxygen (40 mh^−1^). When third true leaves were developed, the following treatments were applied: the seedlings cultured in half-strength Hoagland nutrient solution with or without 75 mM NaCl. The leaves were sprayed with 8 mM Put, 0.1 mM glutathione ethyl ester (GSHmee), 2 mM D-arginine (D-Arg, an inhibitor of putrescine synthesis), 10 mM aminoguanidine hydrochloride (AG, ketamine oxidase inhibitor), 2 mM,1,8-octanediamine (1,8-DO, polyamine oxidase inhibitor), 1 mM diphenyl iodide chloride (DPI, NADPH oxidase inhibitor), 5 mM salicylyl hydroxamic acid (SHAM), 0.1 mM 1-chloro-2,4-dinitrobenzene (CNDB, GSH scavenger). The concentrations of the above treatments were determined according to the preliminary-experiments. Before each treatment, the inhibitors were sprayed in leaves 12 and 6 h in advance and placed in incubator with light intensity of 1000 μmol·m^−2^·S^−1^ for 6 h to ensure the completely opened the stomata. Leaves were sprayed with Put every day at 5.00 p.m. The second functional leaf was collected and quickly put into liquid nitrogen and stored in −80°C for subsequent analysis.

### Determination of growth indexes, photosynthetic parameters and leaf surface temperature

The plant height, stem diameter and fresh weight of the above-ground of cucumber seedlings were measured with a ruler, digital caliper and electronic balance, respectively. To determine the dry weight seedlings were dried at 100°C for 48 h. Net photosynthetic rate (P_*n*_), stomatal conductance (G_*s*_) and intercellular CO_2_ concentration (C_*i*_) were measured using a portable photosynthesis system (Li-6400, USA) at 9:00 to 11:00 a.m. The leaf chamber temperature (25 ± 1)°C, the light intensity 1000 μmol·m^−2^·s^−1^, CO_2_ concentration (380 ± 10) μmol·L^−1^, and the relative humidity 60%-70% were maintained during the data measurement. The stomatal limitation was determined with this formula: Ls=(C_*a*_-C_*i*_)/C_*a*_*100%, where, C_*a*_ is the atmospheric CO_2_ concentration. The second functional leaf was selected and measured by infrared thermal imager (Testo 8751, Testo, Schwarzwald, Germany), and the image was processed by Testo Comfort Software Basic 5.0 (Testo, Schwarzwald, Germany).

### Determination of stomatal index

The area near the central vein of the second functional leaf of the seedling was finely cut. Gently scrape off the mesophyll on the transparent tape with flat tweezers and water. The adhesive tape with lower skin is made into pieces. The slides were observed with a fluorescence microscope (Leica DM2500, Solms, Germany) at 10×20 times, and photos were taken by Leica application suite V3 (Leica, Solms, Germany). The length and width of pores were measured by ImageJ software (Rawak Software Inc., Stuttgart, Germany). At least 40 stomas were randomly selected for each treatment.

### Determination of H_2_O_2_ and reduced glutathione (GSH) content in guard cells

According to An *et al*. (2016), the content of H_2_O_2_ was determined. The changes of H_2_O_2_ content in guard cells were observed by 2, 7-dichlo rodihydrofluorescein diacetate (H_2_DCF-DA; Sigma, USA). The leaves were cut and placed in the buffer solution containing of 50 μM H_2_DCF-DA (50 mM KCl, 0.1 mM CaCl_2_, and 10 mM MES, pH 6.1) for 20 minutes (dark). After washing three times with non-dye buffer solution, immediately strong adhesive tape was used to stick to the back of the blade. Scrape off the mesophyll from the front of the blade with flat end tweezers, and observed the adhesive tape with lower epidermis. The content of GSH in stomata was determined by monochlorobimane (MCB, Sigma, USA) with the treatment method of Akter *et al*. (2010). At room temperature, the epidermis of the second functional leaf near the main vein of the seedling was removed and placed for staining in a buffer solution containing of 100 μm MCB for 2 h. In guard cells, MCB and GSH form fluorescent glutathione S-bimane (GSB). The images were observed by laser confocal microscope with the excitation 488/405 nm, 481-549/400-495 nm excitation signals were collected to observe DCF and GSB; Zeiss LSM 800 META. Chloroplast auto-fluorescence was used as control (561nm excitation, 562-700nm excitation signal). The image was analyzed with ImageJ software (Rawak Software Inc., Stuttgart, Germany), and each process was repeated three times. The image obtained by CLSM with Carl Zeiss LSM software (ZEN 2018) using a 20x magnification objective.

### Determination of H_2_O_2_ and reduced glutathione content in leaves

The H_2_O_2_ content in leaves was estimated according to the method of Alexieva *et al*. (2001). For the calculation of GSH, we followed the Rao *et al*. (1995) method.

### Determination of polyamine content

The content of polyamine in leaves was determined according to Liu *et al*. (2002). 0.3g fresh leaves sample was homogenized in 1.6 mL of precooled 5% perchloric acid (PCA) followed by kept in an ice bath for 1 h and then centrifuged at 12000 rpm at 4°C for 20 min. The supernatant was collected to estimate the contents of free and conjugated PAs, and the resulting precipitate was used to assay bound PAs. For estimation of free and conjugated PAs, 0.7 mL of supernatant was transferred into a new tube, and 1.4 mL of 2 mol L^−1^ NaOH and 15 μL benzoyl chloride were added with this supernatant, and the mixers were vortexed for 20 s followed by incubation at 37°C for 30 min. After incubation, 2 mL saturated NaCl and 2 mL precooled ether were added in above mixers for oscillation, and the whole solution was gently shaken followed by centrifugation at 3000 rpm for 5min. Later, 1mL of ether phase solution was dried, and then re-dissolved it with the addition of 100 μL methanol (60% W/V) and 20 μL solution was injected to determine the free and conjugated PAs. For estimation of bound PAs, the remaining precipitation was washed again with 5% PCA, followed by centrifugation at 1000 rpm for 5min to remove the supernatant, and repeated it for 4 times. To suspension the precipitation, added 5 mL of 6 mol L^−1^ HCl and sealed in an amperometric flask, hydrolyzed followed by incubation the solution at 110°C for 18 h. After incubation, the solution was filtered, evaporated to dryness at 70°C, and the residue was re-suspended and dissolved with 1.6 mL of 5% PCA, and centrifuged for 20 min at 12000 rpm at 4°C for 1h. The sample preparation of bound PAs quantification, the above-mentioned procedure was used. The whole PAs determination was carried out by ultra-performance liquid chromatography (Ultimate 3000, Thermo Scientific, San Jose, CA, USA).

### Determination of abscisic acid content

For ABA extraction Dobrev and Kaminek (2002) protocol and for purification Albacete *et al*. (2008) method was used. Briefly, 0.3 g of leaf was homogenized in 3 mL precooled 50% chromatographic methanol (v/v), and the extracted was incubation at 4°C for 12h followed by centrifugation at 10000 rpm at 4°C for 10 min, and then, the supernatant was stored at 4°C. Again, 2 mL of precooled 80% methanol was added to the residue, extracted at 4°C for 12 h, and centrifuged at 10000 rpm at 4°C for 10 min. After that, 2mL of precooled 100% methanol was added to the residue and extracted for 12 h, and centrifuged as the above condition. Finally, all the extracts were collected and combined, and PVPP was added into the extract at the rate of 0.2 g^−1^·FW to adsorb phenols and pigments. After shaking at 4°C for 60 min, centrifuged the mixed solution like the above condition. The supernatant was passed slowly through the C18 column, collected in a centrifugal tube, and then kept into freeze-drying machine. Thereafter, 2-5 mL of 50% methanol was added in supernatant for dissolution, passed through 0.22 μm organic phase ultrafiltration membrane for determination of ABA. Chromatographic conditions: Hypersil ODS C18 column (250mm × 4.0mm, 5μm). The mobile phase: methanol and ultrapure water (0.5% glacial acetic acid added). For determination of the quantitative value of ABA external standard calibration curve method was used, and each sample repeated three times.

### Determination of enzyme activity

To determine the different enzyme activities, firstly, 0.5 g fresh sample was ground in 1.6 mL phosphoric acid buffer (0.1mol L^−1^, pH 6.5) followed by centrifugation at 4°C for 20 min at 10000 rpm, and the supernatant was collected for amine oxidase extraction. The 3 mL reaction system consisted of 0.1 M phosphate buffer, 2 mM N, N-dimethylaniline, 0.5 mM 4-aminoantipyrine, 0.1 mL of 250 U mL peroxidase solution and 0.2 mL extraction solution. The OD value was measured at 550 nm at 0 min and 30 min interval using an UV-visible spectrophotometer (UV-2600, Unico, Shanghai, China). The activity of DAO was measured when Put was used to start the reaction (Rea *et al*., 2004), and the PAO activity was measured by using Spm to start the reaction (Mhaske *et al*., 2013).

The NOX activity was determined using plant NADPH oxidase ELISA Kit (MEIMIAN, China) with the manufacturer’s instructions. The CWPOD activity was measured according to Lin and Kao, (2001) using protocol.

### Total RNA extraction and real-time fluorescent quantitative PCR (qRT-PCR) analysis

The total RNA content in leaves was extracted using the Total RNAsimple Kit (Tiangen Biotech Co., Ltd, Beijing, China) in accordance with manufacturer’s guidelines. 1μg RNA was reverse transcribed into single-stranded cDNA using the SuperScript First-strand Synthesis Kit (Takara Bio Inc., Tokyo, Japan). Gene-specific primers were designed by Beacon Designer 7.9 software (Premier Biosoft International, CA, USA) based on nucleotide sequences retrieved from a search of the National Center for Biotechnology Information databases (NCBI) and the Cucumber Genome Database (http://cucumber.genomics.org.cn). StepOnePlus™ fluorescence quantitative PCR (Applied Biosystems) and SYBR ^®^ Premix Ex Taq™ II kit (Takara) were used for qRT-PCR target fragment. The quantification of qRT-PCR was repeated three times. The thermal cycling conditions were as follows: 95°C for 5 min, followed by 40 cycles at 95°C for 15 s, 60°C for 1 min, and finally extended at 95°C for 15 s. The relative expression of genes was analyzed according to Jahan et al. (2019). The relative mRNA expression level was calibrated with *actin* (internal standard gene) and compared with the control group.

### Statistical analysis

At least five (5) seedlings per treatment were used to collect samples, and all measurements were repeated at least 3 times. Each experiment contains at least three biological replicates. All data were statistically analyzed by SPSS 17.0 software (SPSS Inc., Chicago, IL, USA), Comparisons of means were performed using Duncan’s multiple range test at P ≤ 0.05 level of significance. The stomatal aperture data were collected from three independent experiments.

## Results

### 1. Effect of exogenous Put on growth of cucumber seedlings under salt stress

As shown in Fig. 1, after 7 days of treatment, there was no significant differences in the growth of cucumber seedlings treated with Put alone. Exogenous Put significantly increased plant height, shoot fresh and dry weight under salt stress. Conversely, compared with salt stress, these growth parameters increased by 36.58%, 8.12% and 17.75%, respectively. However, the stem diameter did not change significantly (P<0.05). These results indicated that exogenous Put could alleviate salt-induced growth inhibition of cucumber seedling by increasing biomass accumulation under salt stress.

**Fig. 1.**
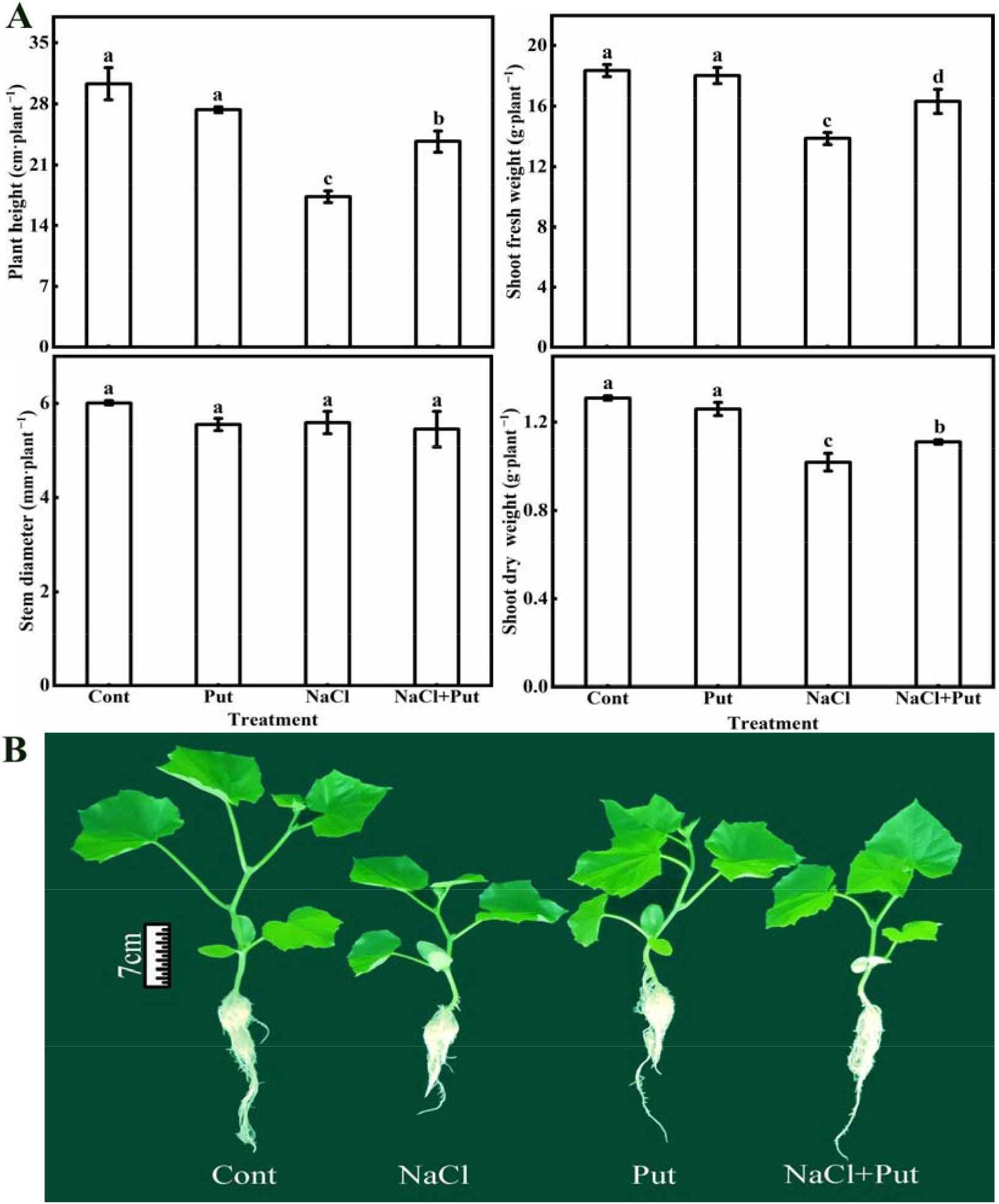
Effect of exogenous put on Cucumber Seedling Growth under salt stress. Cont: The seedlings were cultured in normal nutrient solution, and the leaves were evenly sprayed with distilled water. Put (8mmol L^−1^): The seedlings were cultured in normal nutrient solution and sprayed with put evenly on the leaves. NaCl: The seedlings were cultured in 75mmol L^−1^ NaCl nutrient solution, and the leaves were evenly sprayed with distilled water; NaCl+Put: The seedlings were cultured in 75mmol L^−1^ NaCl solution, and Put was sprayed evenly on the leaves; A: After 7 days, exogenous Put alleviated the growth inhibition of cucumber seedlings under salt stress. B: After 7 days, the effect of exogenous Put on the phenotype of cucumber seedlings under salt stress. Each data represents the mean±SE of three independent experiments, the vertical bar represents SE (n=3), and different letters indicate the significant difference when P<0.05.

### 2. Effects of exogenous Put on photosynthesis and stomatal characteristics of cucumber seedlings under salt stress

One day (1) day after salt treatment, photosynthesis parameter namely: P_*n*_ and G_*s*_ were significantly reduced by 63.7% and 65%, respectively as compared with the control group (Fig. 2). On the other hand, foliar sprayed with Put in salt-treated seedlings increased these components by 90.89% and 71.43%, respectively than those the salt treatment alone. Interestingly, with the increases of treatment time, the improvement effect of exogenous Put on P_*n*_ and G_*s*_ was gradually obvious. After 7 days of salt treatment, C_*i*_ value was higher than that of the corresponding control group for the first time, and Ls was lower than that of the control group, indicating that the salt stress was mainly caused stomatal limitation and changed into non-stomatal limitation. Exogenous Put could prolong the transformation time under salt stress environment (P<0.05). As shown in Figure S1, one day after Put treatment, stomatal morphology and leaf surface temperature were significantly improved under salt stress. Since the photosynthetic parameters changed significantly one day after treatment, we focused our eyes on this period. The results showed that the decreased of P_*n*_ was mainly due to stomatal limitation. Exogenous Put could relieve stomatal limitation by improving the cooperation among G_*s*_, C_*i*_ and P_*n*_ under salt stress.

**Fig. 2.**
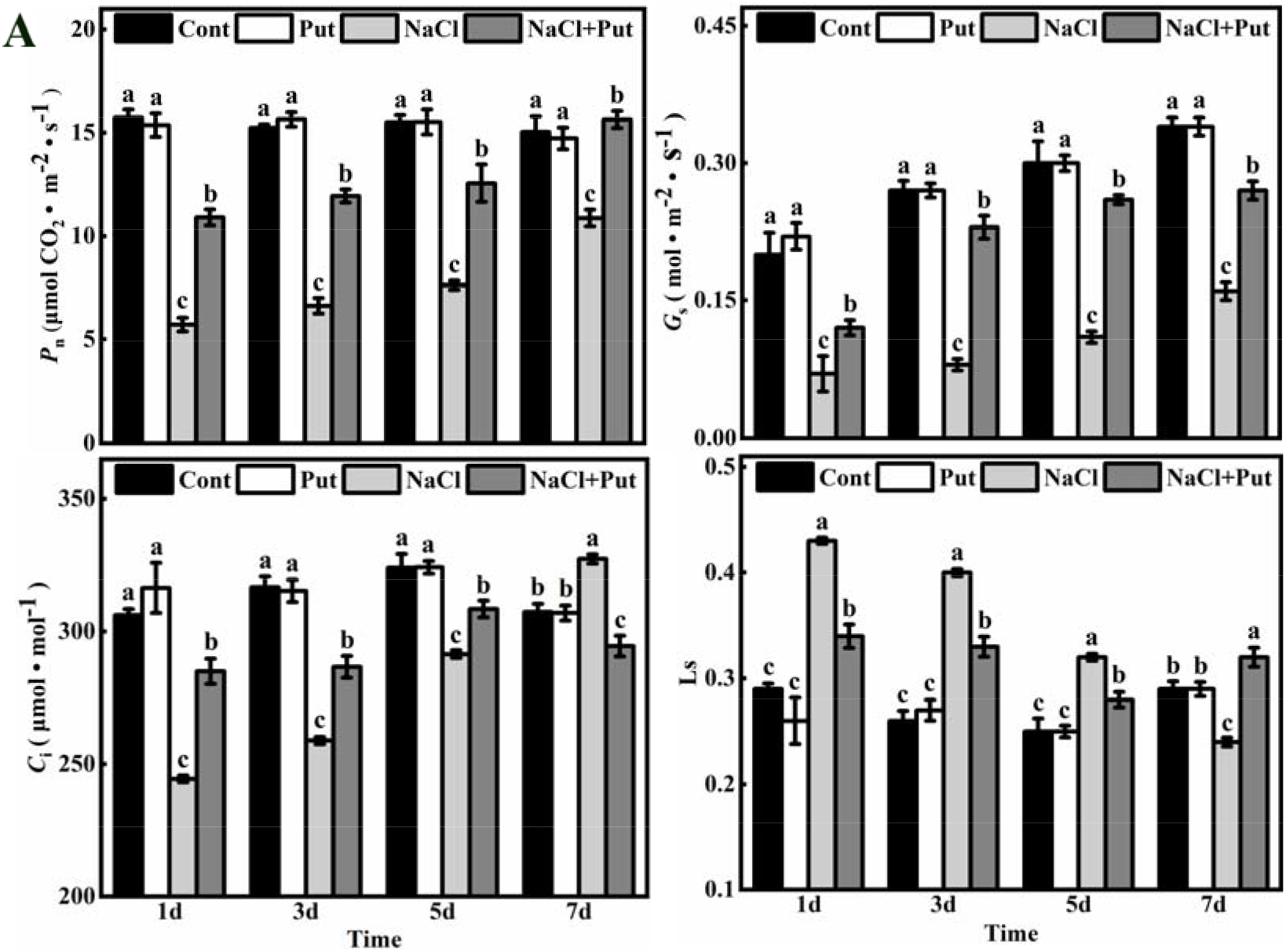
Effects of exogenous Put on P_*n*_, G_*s*_, C_*i*_ Ls of cucumber under salt stress. A: After 1, 3, 5 and 7 days, the effects of exogenous Put on P_*n*_ (net photosynthetic rate), G_*s*_ (stomatal conductance), C*_i_*, (intercellular carbon dioxide concentration) and Ls (stomatal limitation) of cucumber leaves under salt stress were studied. Each data represents the mean ± SE of three independent experiments, the vertical bar represents SE (n=3), and different letters indicate the significant difference when P<0.05.

### 3. Effects of different treatments on the content and distribution of endogenous polyamines

The PAs content of Put treated in salt-stressed seedlings showed a trend of increasing first and then decreasing, reaching the highest level after 6 h (Fig. 3A). To determine the response of Put, Spd and Spm, we analyzed their distribution in total PAs. As shown in Fig. 3A, the contents of these three PAs in the control group were changed smoothly and kept at the same order of magnitude. The content of total polyamines increased rapidly to the peak and maintained this level only in the Put treatment seedlings and it was reached in the highest position and recorded as 998.48 nmol·g^−1^ FW (P<0.05). After salt treatment, only Put content showed a trend of increasing first and then decreasing, reaching the peak at 9 h, accounting for 64.40% of the total polyamine content. After the seedlings were treated with Put under salt stress, the content of Put, Spd and Spm showed a trend of total polyamines, reaching the peak after 6 h, accounting for 65.75%, 22.30% and 11.95% of total polyamines respectively (P<0.05). The results showed that the content of PAs was changed by Put. To get more insights about this, we quantified the expression pattern of Put synthetase genes *ADC 1, 2, 3, 4*. In salt-stressed with or without Put treatment seedlings, *ADC2, 3* showed a trend of increasing first and then decreasing. In respect to the control group, *ADC2* increased 1.76, 1.98 folds and *ADC3* increased 2.52, 2.78times, respectively. It should be noted that the expression levels of *ADC2* and *ADC3* genes were significantly increased by exogenous Put under salt stress (P<0.05). As shown in Figure S3, under each treatment, the PAs content and *ADC2, 3* changed significantly. The results showed that exogenous Put could increase the content of polyamines in leaves by induced and non-induced ways. Since *ADC1* and *ADC4* changed significantly after 1 day, which was beyond the time period of our study, no further analysis was needed.

**Fig. 3.**
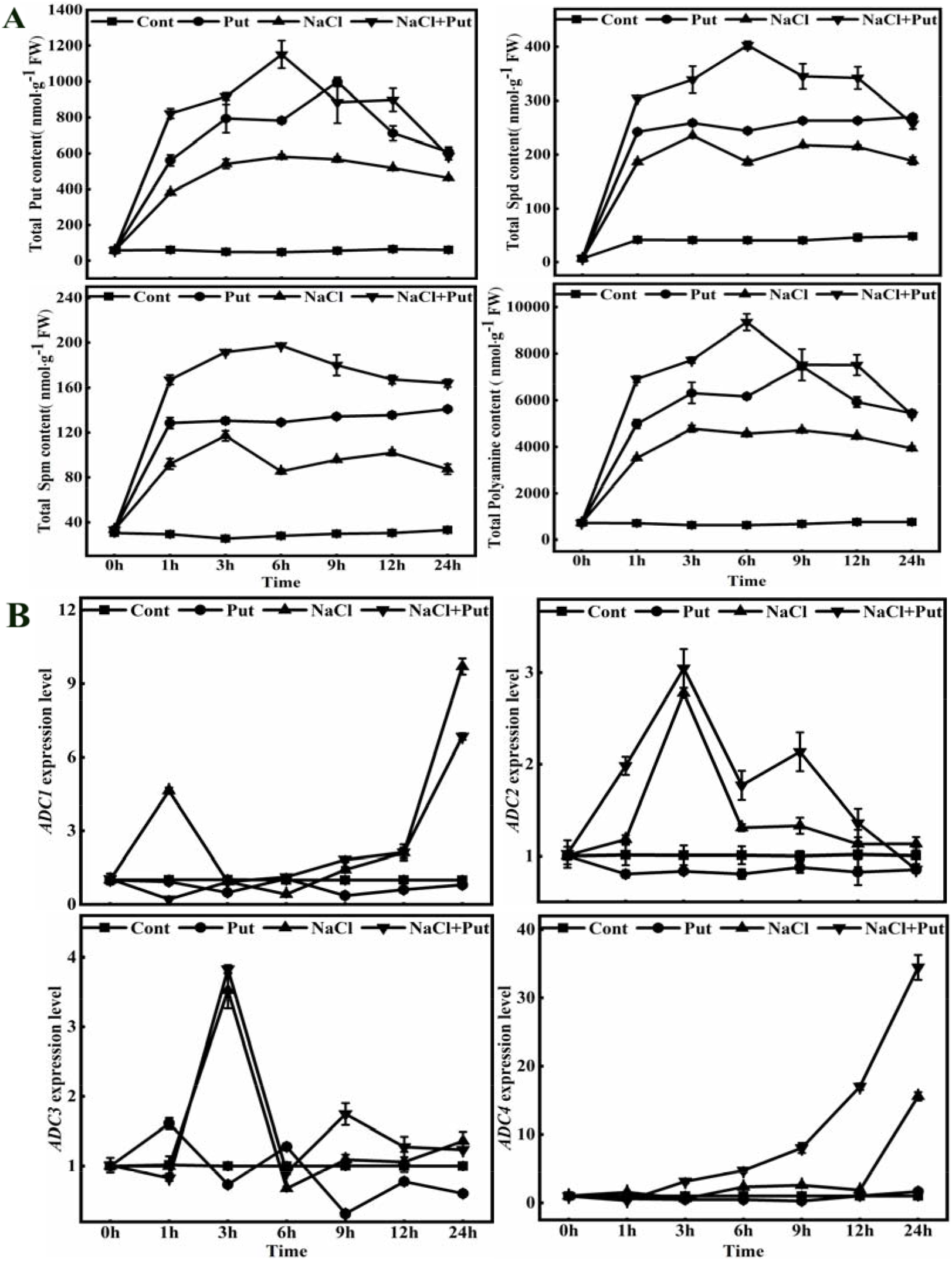
Effects of different treatments on PAs content distribution, total PAs content and gene expression of Put synthesis key enzyme in cucumber leaves within 24hours. A: Effects of different treatments on the contents of three main PAs (Putrescine, Put, Spermidine, Spd, Spermine, Spm) and total PAs in cucumber leaves within 24hours. B: The changes of key genes of put synthetase in 24 hours under different treatment conditions. Each data represents the mean ± SE of three independent experiments, the vertical bar represents SE (n=3), and different letters indicate the significant difference when P<0.05.

### 4. Regulatory function of Put on stomatal-aperture is related to H_2_O_2_ under salt stress

There was no significant change in stomatal aperture in solely Put treatment seedling leaves (Fig. 4A). Conversely, in salt-stressed plants, the stomatal aperture decreased continuously, and reached a significant level after 3 h, and decreased the lowest level after 24 h. However, exogenous Put could effectively alleviate the decrease of stomatal-aperture under salt stress, and the effect was the best after 6 h, and remained stable. As shown in Fig. 4C, the peak appeared only 1 h after Put treatment, which increased by 44.04% compared with the corresponding control group and decreased to the control level after 3 h. The content of H_2_O_2_ was increased first and then decreased both in salt with or without Put treatment, and these two peaks appeared after 1h and 6 h, respectively. As compared with only salt treatment, the content of H_2_O_2_ was increased by 19.52% and 28.14%, respectively in the salt with Put treated plants. Compared with the control, the H_2_O_2_ accumulation in the salt treatment group increased by 71.56% and 234.78%, respectively. As shown in Fig. 4B, the changing trend of DCF fluorescence intensity in stomata is similar to that of H_2_O_2_ content in leaves. These results suggest that H_2_O_2_ may participate in the regulation process in the form of signal molecules.

**Fig. 4.**
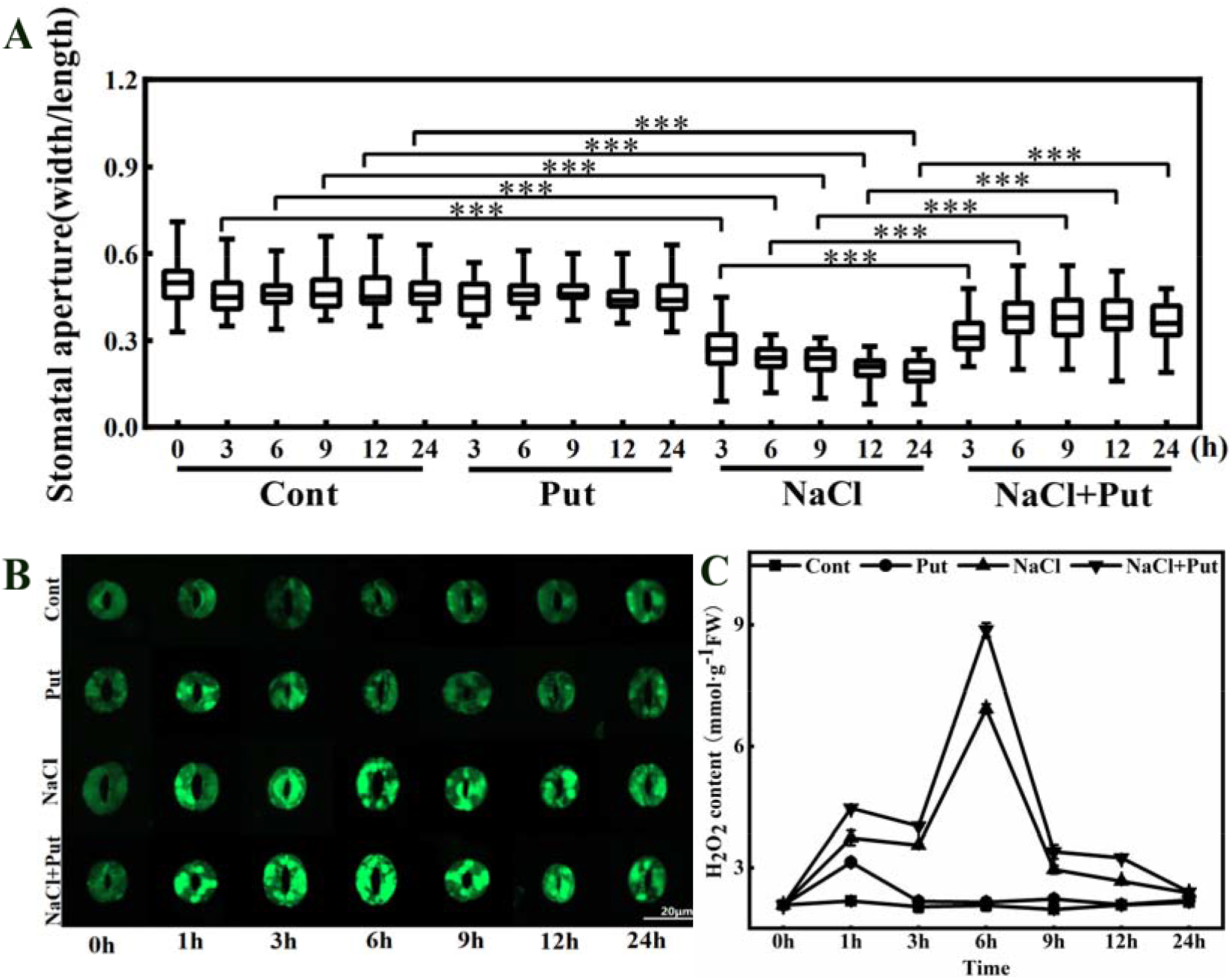
Effects of exogenous Put on stomatal aperture and H_2_O_2_ content under salt stress for 24hours. A: The effect of each treatment on the stomatal aperture. Data were obtained from three independent experiments, ***P<0.001 (n=120 for each treatment), as analyzed by one-way ANOVA followed by a Tukey Kramer post hoc test. B, C: Effects of different treatments on H_2_O_2_ content in guard cells and leaves. The fluorescence of DCF was green with a scale of 20μm. Each data represents the mean ± SE of three independent experiments, the vertical bar represents SE (n=3), and different letters indicate the significant difference when P<0.05.

### 5. The alleviating effect of Put on stomata under salt stress is based on the regulation of ABA content by H_2_O_2_

Although the content of endogenous PAs and H_2_O_2_ reached the peak after 6 h, the stomatal-aperture changed significantly after 3 h. The results showed that the signal of stomatal changed was completed at/or before 3 h. Therefore, we have chosen 3 h time point for further study. As shown in Fig. 5A, C, D, the stomatal-aperture of salt with Put treatment decreased to the level of salt treatment compared with that before application of D-Arg and DMTU and the fluorescence intensity of DCF in stomata decreased by 0.2, 0.77 times, respectively. The results employed that Put had a certain specificity on stomatal-aperture regulation under salt stress, and endogenous PAs played a regulatory role in the upstream of H_2_O_2_. The ABA as a key signal substance in stomatal regulation under salt stress has been deeply studied. In order to determine whether ABA is involved in the regulation process, we determined the endogenous content of ABA in each treatment condition. As shown in Fig. 5B, Put alone had no significant effect on ABA content. Exogenous application of Put markedly inhibited the increase of ABA content under salt treatment, and it was 0.79 folds lower than salt-stressed leaf. The ABA content of salt with Put treatment decreased by 6.36 times compared with that before application of DMTU, but the stomatal aperture increased to a significant level. Supplemental application of H_2_O_2_, the ABA content decreased by 4.77 times, but the stomatal-aperture is reduced to a very significant level. These results indicate that ABA is involved in the regulation of Put on stomata, and PAs and H_2_O_2_ play a role in the upstream of ABA. In order to further elucidation of the underlying molecular mechanism on how H_2_O_2_ affects ABA, we quantified transcript abundance of the four genes that induce ABA synthesis gene *RD29B*, ABA signal transduction key gene *Snrk*, and ABA signal pathway negative regulation gene *ABI* under each treatment condition (Fig. 5E). Interestingly, only ABA synthesis gene *NCED* showed significant effect on the different treatments and it was consistent with the changes in endogenous ABA content.

**Fig. 5.**
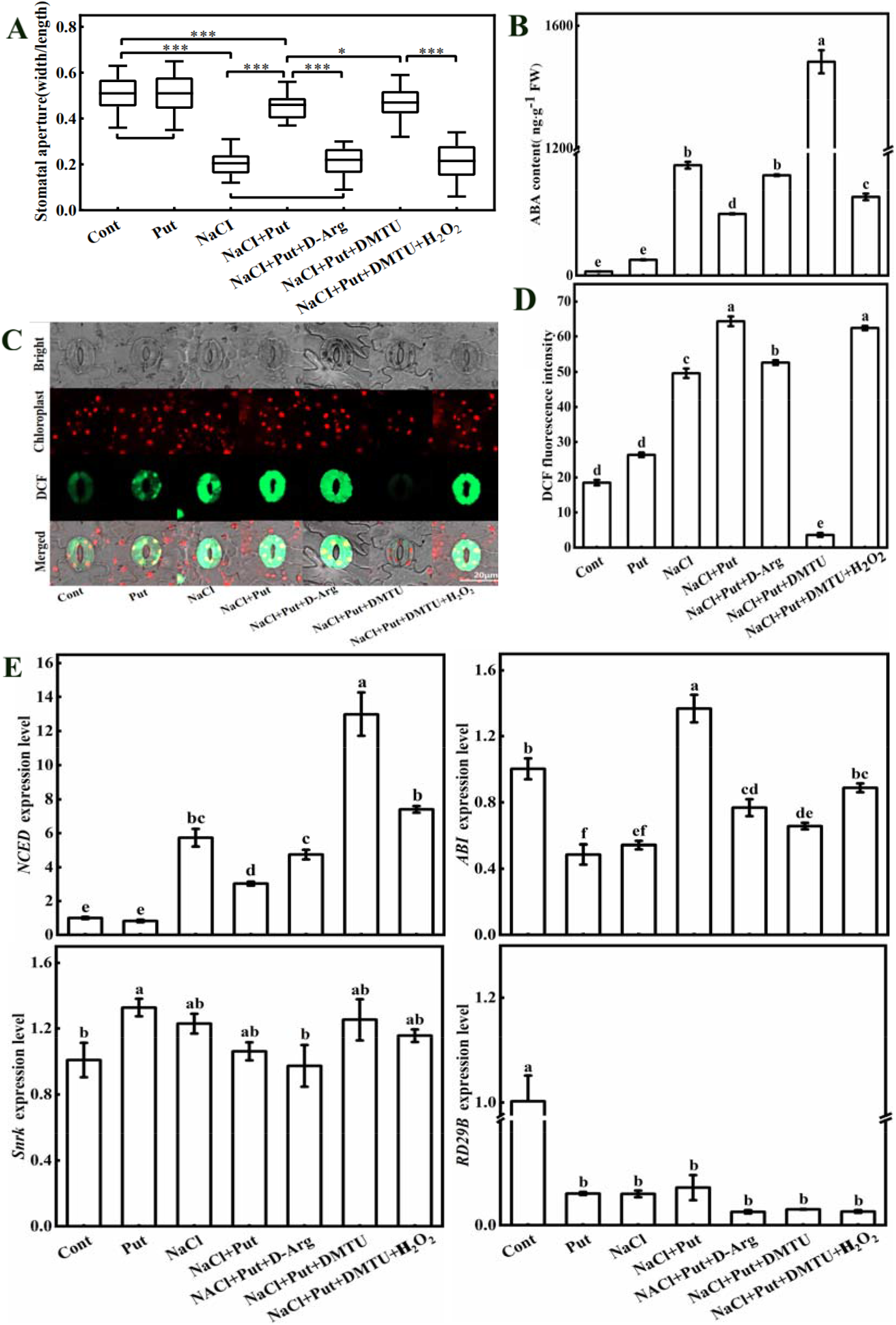
H_2_O_2_ and ABA participate in the alleviation mechanism of exogenous Put on stomatal closure of cucumber leaves under salt stress. A: The effect of each treatment on the stomatal aperture, Among them, 0.2 mM H_2_O_2_ of NaCl + put + DMTU + H_2_O_2_ treatment was sprayed on the leaves one hour before sampling. Data were obtained from three independent experiments, *P<0.05, ***P<0.001 (n=120 for each treatment), as analyzed by one-way ANOVA followed by a Tukey Kramer post hoc test. B: Effects of different treatments on ABA content in cucumber leaves. C, D: The effects of different treatments on H_2_O_2_ content and DCF fluorescence intensity in guard cells were studied, the fluorescence of DCF was green and the spontaneous fluorescence of chloroplast was red with a scale of 20 μm. E: The effect of each treatment on ABA synthase gene expression. Each data represents the mean ± SE of three independent experiments, the vertical bar represents SE (n=3), and different letters indicate the significant difference when P<0.05.

### 6. H_2_O_2_ effect on the ABA content through the catalysis of DAO, PAO, and NOX

Based on the main pathways of H_2_O_2_ production in plants, we hypothesized that H_2_O_2_ might be catalyzed by DAO, PAO, CWPOD and NOX. As displayed in Fig. 6A, C, D, the stomatal-aperture and DCF fluorescence intensity of salt with Put treatment were not affected by SHAM pre-treatment. The stomatal-aperture of salt with Put treatment decreased to a very significant level than without DPI treatment. At the same time, the fluorescence intensity of DCF was reduced by 13.25%. Compared with SHAM treatment, the stomatal diameter of DPI treatment decreased to a significant level, and DCF fluorescence intensity decreased by 12.54%. In response to pre-treatment with DPI, the stomatal-aperture of AG treatment was significantly reduced, and the fluorescence intensity of DCF decreased by 38.12%. The stomatal-aperture of 1,8-DO treatment was lower than that of DPI treatment, but higher than that of AG treatment, whereas, the DCF fluorescence intensity decreased by 15.9% and increased by 35.90% respectively. Further, we verified the inhibitory effects of these four kinds of inhibitors and as shown in Figure S3, these inhibitors have a significant inhibitory effect on their corresponding enzyme activity as well as their encoding gene expression. These results indicate that H_2_O_2_ plays a regulatory role mainly through the maintaining of DAO, PAO and NOX catalysis. However, there is no significant involved was found of CWPOD for the regulation of ABA content. As indicated in Fig. 6B, E, the endogenous ABA content and the transcript abundance of *NCED* in salt-stressed with Put treatment increased by 25.96% and 38.86%, respectively than that of the prior treatment of SHAM. In respect to the pre-treatment of SHAM, the ABA content and mRNA level of *NCED* in DPI treatment increased by 29.68% and 16.09%, respectively. Similarly, the ABA content and *NCED* expression of AG treatment increased by 125.16% and 80.19% respectively than that of SHAM treated plants and compared with the prior application of DPI, it increased by 73.65% and 56.1 2%, respectively. In addition, in contrast with the pre-treatment of SHAM/DPI/AG, the endogenous ABA level and *NCED* gene expression of 1,8-DO treatment increased by 148.37/73.65/10.28% and 64.22/56.12/9.4%, respectively. The gene expression of *RD29B, Snrk* and *ABI* did not show regular changes under various treatment combinations, so no further analysis was conducted. These results suggest that H_2_O_2_ is mainly catalyzed by DAO, PAO and NOX, and H_2_O_2_ plays a key regulatory role in the upstream of ABA.

**Fig. 6.**
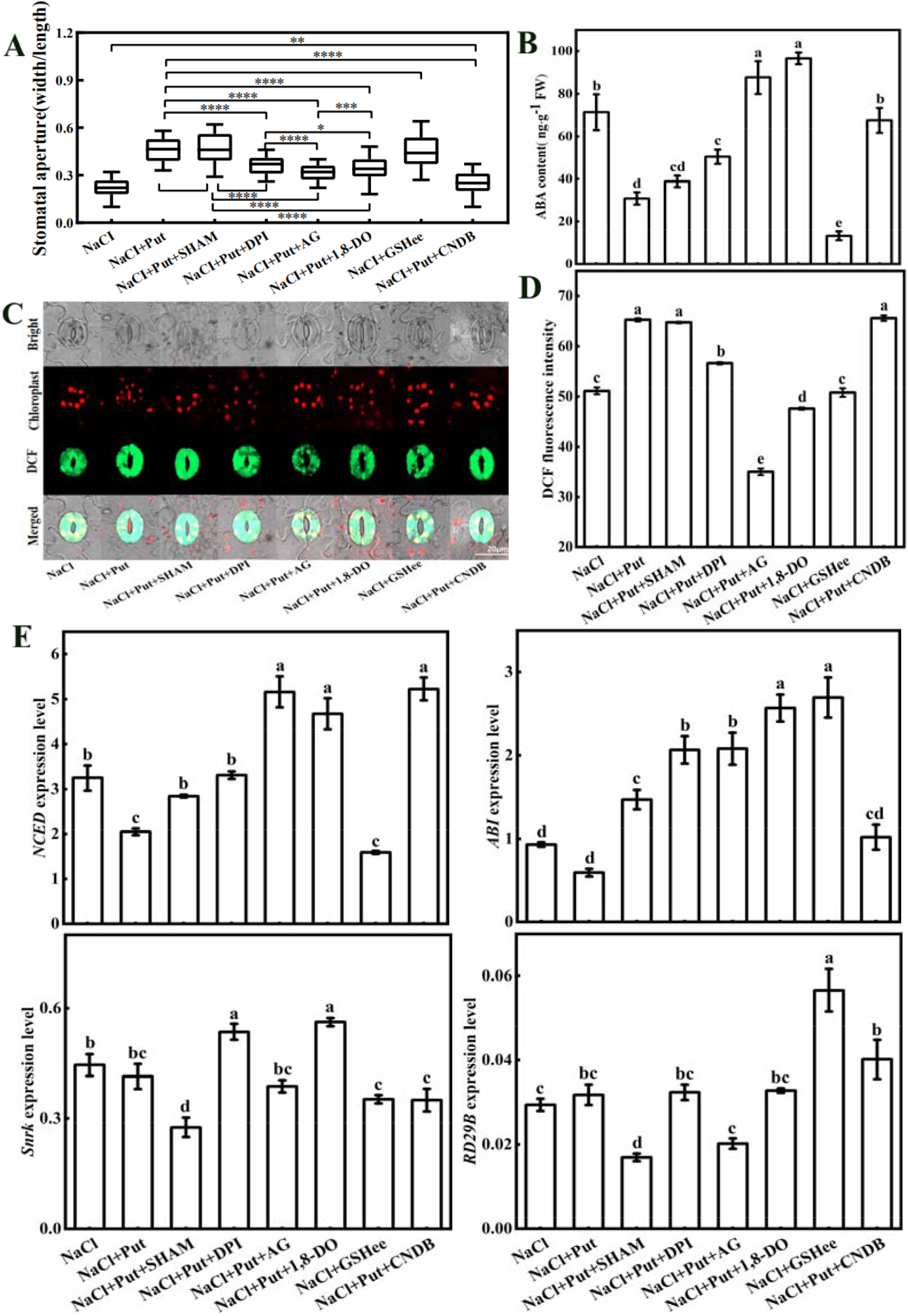
The effects of four enzyme inhibitors (DAO, PAO, NOX and CWPOD) on stomatal aperture, H_2_O_2_ accumulation, ABA content and *NCED* expression in guard cells. A: Effects of 10 mM SHAM, 0.1 mM DPI, 10 mM AG and 2 mM 1,8-DO pretreatment on stomatal aperture under different treatments were studied. Data were obtained from three independent experiments, *P<0.05,**P<0.01,***P<0.001 (n=120 for each treatment), as analyzed by one-way ANOVA followed by a Tukey Kramer post hoc test. B: Effects of pre application of inhibitors and scavengers on ABA content under different treatment conditions were studied. C, D: Effects of pre application of inhibitors and scavengers on H_2_O_2_ content and DCF fluorescence intensity in guard cells under different treatment conditions were studied. The fluorescence of DCF was green and the spontaneous fluorescence of chloroplast was red with a scale of 20 μm. E: Effects of pre application of inhibitors and scavengers on expression level of *NCED* of ABA synthase gene under different treatment conditions were studied. Each data represents the mean ± SE of three independent experiments, the vertical bar represents SE (n=3), and different letters indicate the significant difference when P<0.05.

### 7. GSH is downstream of H_2_O_2_, which is involved in the alleviation mechanism of exogenous Put on stomata of cucumber leaves under salt stress

The above results have shown that GSH induced by H_2_O_2_ and has a negative effect on ABA-mediating stomatal closure. So we assumed that GSH might play a role in this process. The GSH content did not change significantly in solely Put treated plants. However, compared with the control group, the fluorescence intensity of GSB increased by 99.35% (Fig. 7A, B, C). The GSB fluorescence intensity and GSH content of salt-stressed plants were increased by 186.49% and 84.84%, respectively than that of its corresponding control group. In respect to salt treatment, GSB fluorescence intensity and GSH content of salt with Put treatments were increased by 46.7% and 68.85%, respectively. However, the GSB fluorescence intensity and GSH content of salt with Put treatment decreased by 20.09% and 40.78%, respectively than that of the D-Arg treated plants. Similarly, it decreased by 77.40% and 72.82%, respectively than those of the DMTU pre-treatment plants. Interestingly, after supplementing treatment with H_2_O_2_, it was increased by 199.28% and 56.92%, respectively. To further gain more evidence, we quantified the GSH synthase gene expression, as shown in Fig. 7C, the changes of *GSHS1, 2* could fit the changing trend of GSH under each treatment. These results deduce that H_2_O_2_ regulates GSH content at the transcript level.

**Fig. 7.**
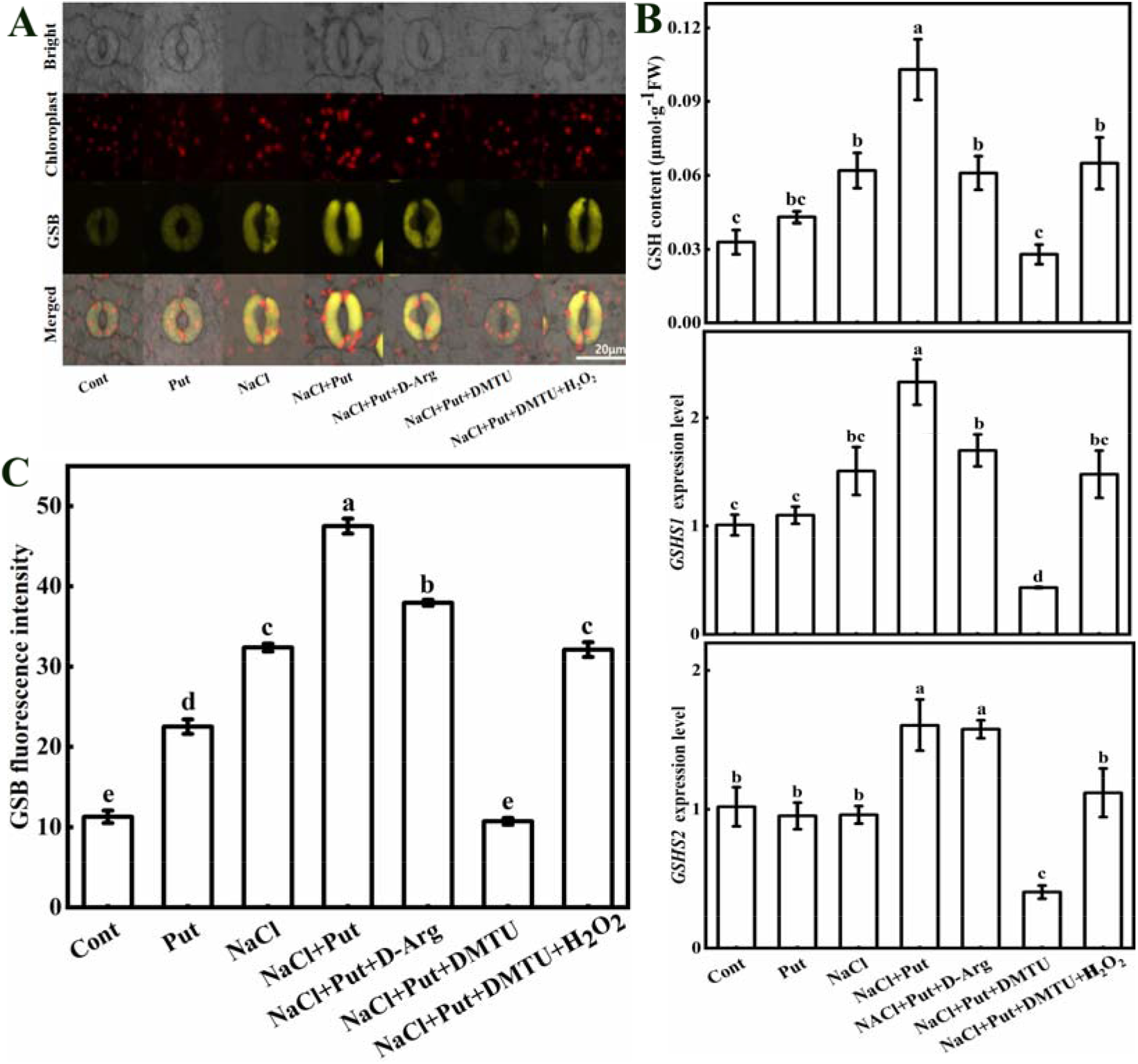
GSH is involved in the mechanism of exogenous Put to alleviate stomatal closure under salt stress. A, C: Effects of different treatments on GSH content and GSB fluorescence intensity in guard cells were studied. The fluorescence of GSB was yellow and the spontaneous fluorescence of chloroplast was red with a scale of 20 μm. B: The effects of different treatments on GSH content and expression of *GSHS1, 2* genes encoding GSH synthase were studied. Each data represents the mean ± SE of three independent experiments, the vertical bar represents SE (n=3), and different letters indicate the significant difference when P<0.05.

### 8. GSH plays a regulatory role on ABA content

In order to further illustrated the relationship between GSH and ABA, we used GSHee and CNDB. As shown in Fig. 6A, the effect of exogenous GSHee on stomatal aperture under salt stress was similar to that of exogenous Put. The stomatal aperture of salt with Put treatment decreased to the level of salt treatment compared with the CNDB pre-treatment plants. Surprisingly, the DCF fluorescence intensity of salt with Put treatment had no significant changes as compared to the salt with GSHee treatment (Fig. 6C, D, 8A, B). However, in respect to salt with or without Put treatments, the GSB fluorescence intensity of salt with GSHee treatment increased by 28.38% and decreased by 5.86%, respectively, and in contrast, the content of GSH increased by 96.77% and decreased by 18.45%, respectively. The DCF fluorescence intensity of salt with Put treatment had no significant effect as compared to prior CNDB treatment. But the GSB fluorescence intensity and GSH content were reduced by 77.97% and 87.38%, respectively compared with the salt with Put treatment. It is suggested that H_2_O_2_ can affect stomatal aperture by affecting GSH content. As shown in Fig. 6B, E, the ABA content and *NCED* expression of salt with Put treatment increased by 118.70% and 154.89% respectively, as compared to prior CNDB treatment, inferring that the regulation of exogenous Put on stomata depends on the change of GSH content under salt stress. As shown in Fig. 8A, B, C, the GSB fluorescence intensity and GSH content of salt with Put treatment reduced by 37.39% and 24.27% respectively, as compared to prior SHAM treatment. Compared with the pre-treatment with SHAM, the GSB fluorescence intensity of DPI treatment increased by 13.38%, GSH content had no significant change. The GSB fluorescence intensity and GSH content of salt with Put treatment reduced by 62.31% and 84.47% respectively, as compared to prior AG treatment. Compared with the pre-treatment of SHAM, the GSB fluorescence intensity and GSH content of AG treatment reduced by 39.80% and 79.49%, respectively. The GSB fluorescence intensity and GSH content of salt with Put treatment reduced by 56.14% and 72.82%, respectively, as compared to prior 1,8-DO treatment. Compared with the pre-treatment of AG, the GSB fluorescence intensity and GSH content of AG treatment reduced by 16.35% and 75.00%, respectively. However, these were decreased by 29.92% and 64.10%, respectively, compared with pre-treatment of SHAM treatment. As shown in Fig. 8C, the changing trend of *GSH1, 2* gene expression was consistent with the above GSH content. These results indicating that H_2_O_2_ produced by DAO, PAO and NOX is involved in the regulation of ABA by GSH.

**Fig. 8.**
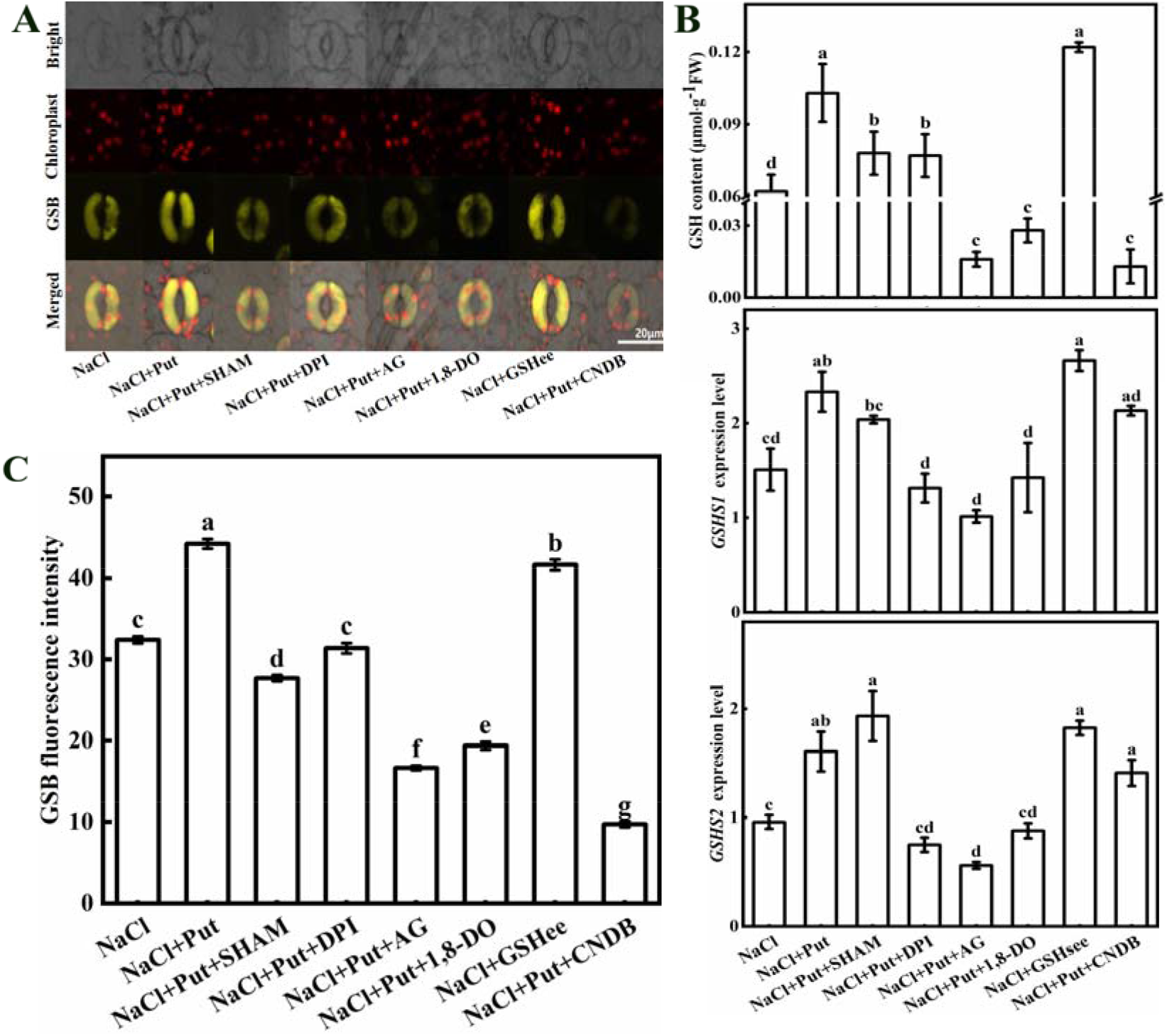
The effects of four enzyme inhibitors (DAO, PAO, NOX, CWPOD) on GSH accumulation in guard cells, GSH content in leaves and expression of key genes *GSHS1, 2* in leaves. A, C: Effects of 0.1mmol L^−1^ CNDB, 0.1mmol L^−1^ GSHmee pretreatment on GSH content and GSB fluorescence intensity in guard cells under different treatments were studied. The fluorescence of GSB was yellow and the spontaneous fluorescence of chloroplast was red with a scale of 20 μm. B: The effect of each treatment on GSH content and expression of *GSHS1, 2* in leaves were studied. Each data represents the mean ± SE of three independent experiments, the vertical bar represents SE (n=3), and different letters indicate the significant difference when P<0.05.

## Discussion

Recently, most of the photosynthesis based research under salt stress has mainly focused on improving the stability of photosynthetic organs and antioxidant capacity (Ioannidis *et al*., 2012; Saha *et al*., 2015). However, very few studies have been emphasized on the mechanism of PAs to enhance photosynthesis by improving stomatal-aperture in the form of the signal during the stomatal limitation period. H_2_O_2_ plays an important role in the signal transduction process of resisting salt stress (Li *et al*., 2015; Niu and Liao, 2016). It is noteworthy that H_2_O_2_ is one of the direct products of oxidative metabolism of PAs, which lays a foundation for the study of polyamine signaling in the stomatal limitation period under salt stress (Pottosin *et al*., 2014). In the present study, the results showed that exogenous Put could effectively regulate the stomatal-aperture closure of cucumber leaves under salt stress through signal transduction pathway.

### Exogenous Put alleviated the growth inhibition of cucumber seedlings by improving G_*s*_ during stomatal limitation period under salt stress

Exogenous application of PAs have been an effective tool to improve crop stress resistance and crop productivity in response to a variety of abiotic stresses (Lee *et al*., 2012; Hao *et al*., 2012). In this study, exogenous Put significantly alleviated the salt-induced growth inhibition effect of cucumber seedlings during the stomatal limitation period under salt stress (Fig. 1), which is consistent with the previous results (Zhang *et al*., 2009). After salt treatment, as shown in Fig. 2A, the changing from stomatal limitation to non-stomatal limitation was observed in this study and this kind of similar changing trend also found by Munns *et al*. (1993). Exogenous Put not only increased P_*n*_ by improving G_*s*_ (Sun et al., 2016), but also delayed the transition of two periods, which is mainly due to the protective effects of Put on photosynthetic organ stability (Ioannidis *et al*., 2012), the effectiveness of loop electron transport chain (Wu *et al*., 2019) and gas exchange persistence (Zhang *et al*., 2009) under salt stress. In contrast, exogenous Put alone does not seem to cause significant changes, which is consistent with the results of Wu *et al*. (2019), suggesting that exogenous Put may be stimulated by other substances.

### Endogenous PAs and H_2_O_2_ inhibited the increase of ABA content by signal transduction

After salt treatment, as shown in Fig. 4A, stomatal aperture continued to decrease (Zhang *et al*., 2020) and reached the lowest level at 24 h. Exogenous Put could effectively alleviate this trend. However, some studies have found that exogenous Put promotes stomatal closure (An *et al*., 2008). This may be occurred due to the difference of species and treatment conditions, so this is the first time, we investigated the positive regulatory effect of exogenous Put on cucumber stomata under salt stress. The PAs and H_2_O_2_ content showed a typical signal change trend with the supplementation of Put under salt stress, and Freitas *et al*. (2018) studied found the similar effect on ethylene signaling. It should be noted that exogenous Put alone increased the content of endogenous PAs, even exceeded than the salt treatment, but had no significant effect on stomatal-aperture, which further indicated that exogenous Put might be needed for modulation of salt effect to play a positive role. As shown in Fig. 2A, Put had the most significant changes compared to Spd and Spm (Yang *et al*., 2007), which may be happened due to the fact that Put is the precursor of Spd and Spm synthesis (Pál *et al*., 2015). By analyzing the key Put synthase genes, it was found that exogenous Put increased *ADC* expression after salt treatment, but exogenous Put alone did not affect the expression of *ADC*, which indicated that the increasing mode of PAs was different under different treatment conditions, and this part still needs to further study.

As mentioned above, even if complete signal changes are observed, the change of stomatal-aperture may occur before the signal change due to the delay of the measurement method and operation process. This kind of similar problem also appears in the study of Xia *et al*. (2014). It has been widely recognized that ABA content increases and stomatal movement is modulated by ABA under salt stress (Hedrich *et al*., 2018). In the current study, exogenous Put decreased ABA transcript abundance under salt stress. In addition, pre-treatment with D-Arg could eliminate the stomatal alleviation through the exogenous Put treatment under salt stress, and ABA content returned to the same level as of salt treatment. Similarly, pre-treatment with DMTU, the stomatal-aperture returned to the identical control level, but ABA content increased rapidly. However, replenishing with H_2_O_2_, the stomatal-aperture decreased to the salt treatment level, and the ABA content was lower than that of the salt treatment level. These findings suggested that H_2_O_2_ plays an important role in regulating ABA content downstream of Put, but it is not unique. There may be other pathways regulating ABA content. It also indicates that H_2_O_2_ may directly regulate stomatal-aperture changes (Roubelakisangelakis *et al*., 2008).

The sources of H_2_O_2_ in plants are diverse (Foyer and Noctor, 2005). As mentioned above, NOX, CWPOD, DAO and PAO are considered as the most important sources of H_2_O_2_ in plants (Simonovicová *et al*., 2004; Rejeb *et al*., 2015). As shown in Fig. 6, except for the CWPOD, the other three enzymes seem to be involved in the regulation of this process. When DAO and PAO activities were inhibited, endogenous ABA content, DCF fluorescence intensity and stomatal aperture were significantly affected. This may be due to the increase of endogenous PAs content after exogenous Put supplementation, which induces the balance mechanism of PAs, and a large number of PAs are decomposed to produce H_2_O_2_. Although some studies have found that H_2_O_2_ after PAs decomposition can promote stomatal closure (An *et al*., 2008), this may differ from species to species and treatment conditions. In conclusion, the role of H_2_O_2_ produced by PAs metabolism is still controversial due to its dual functions, and to clarify these dual effect needs to be further study.

### GSH downstream of H_2_O_2_ regulates ABA content at the transcription level

The GSH is an important regulator of redox homeostasis in plant cells. GSH affects downstream components of ABA signaling pathway and inhibits stomatal closure by regulating redox status in guard cells (Jahan *et al*., 2008; Okuma *et al*., 2011; Akter *et al*., 2013). However, the effect of GSHee pre-treatment as identical as to that of exogenous Put (Fig. 6, 7). Pre-treatment with CNDB eliminated the alleviating effect of Put on salt stress. Different from D-Arg, removing GSH did not affect the fluorescence intensity of DCF, implying that GSH controls stomatal aperture by regulating ABA transcription level. The H_2_O_2_ can increase GSH and GSH/GSSG ratio (An *et al*., 2012; Wimalasekera *et al*., 2011), and GSH is considered to play critical/pivotal role in H_2_O_2_ signaling pathway (Mhandi *et al*., 2010). Pre-treatment with D-Arg reduced the fluorescence intensity of GSB under salt stress combined with Put treatment plants. After removing H_2_O_2_, the fluorescence intensity of GSB decreased and supplementation with H_2_O_2_, it returned to the same level as salt treatment. These results confer that H_2_O_2_ can regulate GSH in a certain concentration range. As shown in Fig. 8, the effect of pre-application of 4 types of H_2_O_2_ inhibitors on the fluorescence intensity of GSH is similar to that of DCF in Fig. 6, further indicating that the H_2_O_2_ regulates GSH is mainly derived from the catalysis of DAO and PAO.

In summary, our research results show that exogenous Put enhanced the photosynthesis activities of cucumber seedlings by delaying stomata closure under salt stress, thereby improving plants tolerance to salt stress. This process mainly mediated by H_2_O_2_ signaling pathway. Under salt stress conditions, the exogenous Put increased endogenous PAs content and enhanced the activities of DAO, PAO, NOX as well as the expression pattern of their encoding genes further up-regulated, and thus t promoting the formation of H_2_O_2_ signal molecules. When the concentration of H_2_O_2_ reached a certain level, the increase of ABA content in cucumber leaves inhibited by inducing the change of GSH content. Finally, the whole mechanism alleviated salt-stress induced stomatal closure in cucumber leaves (Fig. 9). To insights more understanding about stomatal closure under salt stress via H_2_O_2_/ ABA-mediated signaling pathway, our current knowledge may be useful, and we suggest to apply an in-depth molecular approach to elucidate the better mechanism.

**Fig. 9.**
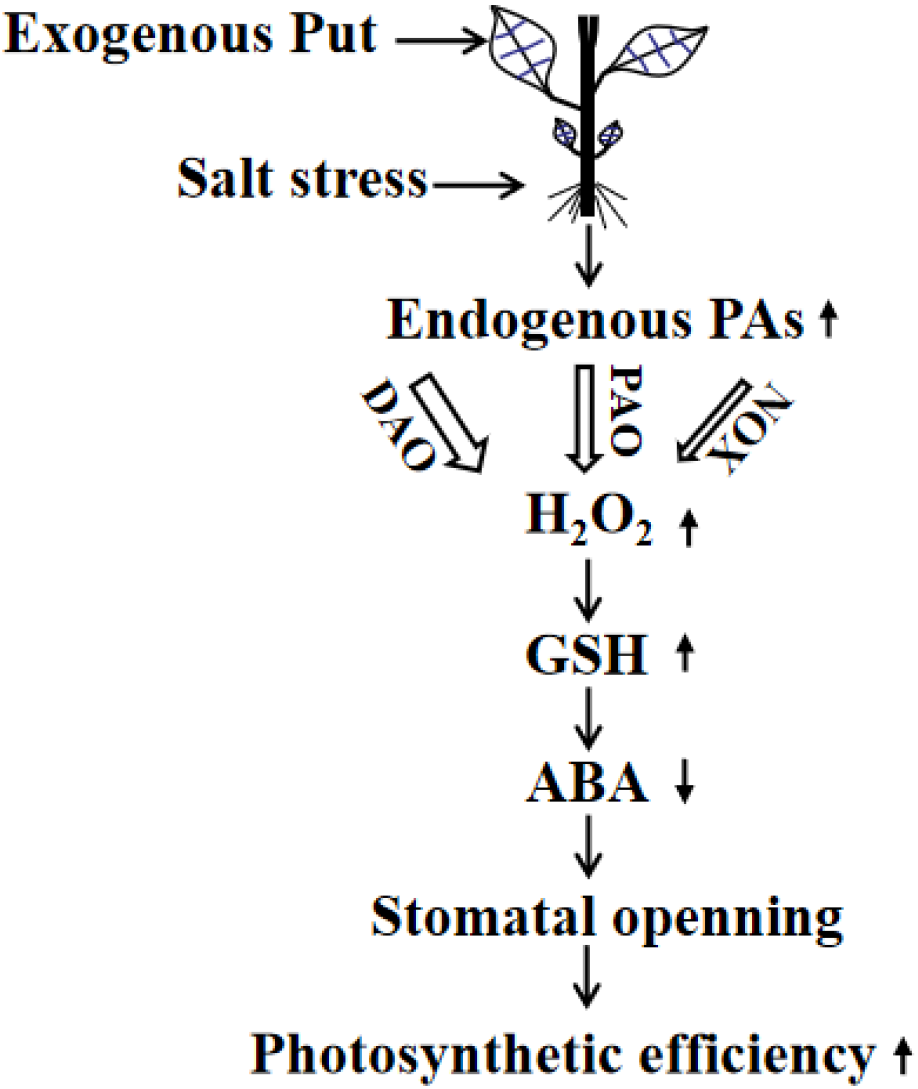
The model diagram of exogenous Put regulating photosynthesis of cucumber leaves by alleviating stomatal closure under salt stress. Put-putrescine, PAs-polyamines, DAO-diamine oxidase,PAO-polyamine oxidase, NOX-nicotinamide adenine dinucleotide phosphate oxidase, H_2_O_2_-hydrogen peroxide, GSH-reduced glutathione, ABA-abscisic acid.

## Data availability statement

The data supporting the findings of this study are available from the corresponding author, (Sheng Shu), upon request.

## Acknowledgements

This work was funded by The National Key Research and Development Program of China (2018YFD1000800), and was sponsored by the China Agriculture Research System (CARS-23-B12).

